# Laboratory evolution of anticipatory gene regulation in *Escherichia coli*

**DOI:** 10.1101/2020.08.20.259606

**Authors:** Anjali Mahilkar, Akshat Mall, Supreet Saini

## Abstract

Environmental cues in an ecological niche are often temporal in nature. For instance, in temperate climates, temperature is higher in daytime compared to during night. In response to these temporal cues, bacteria have been known to exhibit anticipatory regulation, whereby they trigger response to a yet to appear cue, anticipating its actual arrival in the near future. Such an anticipatory response in known to enhance Darwinian fitness, and hence, is likely an important feature of regulatory networks in microorganisms. However, the conditions under which an anticipatory response evolves as an adaptive response are not known. In this work, we develop a quantitative model to study response of a population to two temporal environmental cues, and predict variables which are likely important for evolution of anticipatory regulatory response. We follow this with experimental evolution of *E. coli* in alternating environments of a pentose sugar, rhamnose, and an oxidative stress molecule, paraquat for more than 800 generations. We demonstrate that growth in this cyclical environment leads to evolution of anticipatory regulation, whereby, exposure to rhamnose leads to partial induction of the oxidative stress response regulon. As a result, pre-exposure to rhamnose leads to a greater fitness in paraquat environment. Overall, we show that in niches where environmental stimuli have a cyclical nature, anticipatory regulation can evolve as an adaptive strategy, in a time course of a few hundred generations. This contributes to our understanding of how environment shapes the topology of regulatory networks in an organism.

## Introduction

Specificity is key in biological regulation. Biological systems expend considerable effort towards reducing non-specific substrate binding and reducing crosstalk ^1–4^. However, despite this, regulatory crosstalk between cellular modules is ubiquitous in biology ^5, 6^. One of the forms of specificity and crosstalk in regulation is between transcription factors and their recognition site on the DNA. Although each transcription factor binds its cognate regulatory site on the DNA, significant non-specific binding, and hence regulatory crosstalk, exists ^7, 8^.

A particular form of crosstalk between two functional modules is when gene expression changes upon anticipation of an upcoming environmental shift. For example, in *E. coli*, an increase in temperature elicits response to low oxygen conditions, mimicking how these two environments are sequentially encountered by the bacterium in the mammalian gastrointestinal tract ^9^. Such anticipatory regulation was also demonstrated to occur between the sugars lactose and maltose. In this case, exposure to lactose leads to partial activation of maltose-utilization genes ^10^. In the intestinal gut, exposure to lactose precedes that to maltose. As a result, this partial gene activation provides an adaptive benefit to the organism.

Anticipatory regulation is seen in yeast too. For example, different environmental isolates exhibit varying degrees of glucose-dependent catabolite repression on maltose-utilization genes ^11^. When exposed to a mixture of glucose and maltose, while some strains exhibit strict hierarchical utilization of the two sugars, separated by a diauxic lag, others exhibit minimal lag, and co-utilization of the two sugars. Thus, the regulatory crosstalk between the glucose- and maltose-utilization modules varies among isolates. A similar variation, among natural isolates of yeast, exists in the degree of crosstalk between glucose- and galactose-utilization patterns ^12^. Thus, fine-tuning of regulatory crosstalk between modules, leading to exhibition of anticipatory regulation, is done in accordance with the precise ecological niche of an organism. In a recent study, it was demonstrated that yeast anticipates exhaustion of a primary carbon source, and thus, switches to the secondary source in the environment, even before the primary source is exhausted ^13^.

Anticipatory gene regulation, as a strategy, is widespread in pathogens ^14^. In *Mycobacterium*, the different two-component systems in the bacterium are wired so that activation of one leads to partial activation of the downstream TCSs ^15^. This helps the bacterium respond to an upcoming environmental change faster. In *Salmonella*, flagella is assembled prior to assembly of a *Salmonella* Pathogenicity Island-1 (SPI1)-encoded Type 3 Secretion System (T3SS) in the course of infection ^16^. Regulatory elements in the flagellar cascade are known to activate the SPI1-encoded T3SS genes ^17^. *Burkholderia*, in response to high cell density, anticipates stationary phase stress and triggers anticipatory response ^18^. Fungal pathogens have been shown to elicit anticipatory response for protection against attack from the immune system of the host ^19^.

Thus, repeated, temporal exposure of environmental cues may have led to evolution of regulatory crosstalk between cellular modules. Despite significant evidence that anticipatory gene expression provides an adaptive advantage to an organism, little is known about how anticipatory regulation can evolve in a population.

In this work, we ask the following question: if a population is exposed to two environmental signals, S1 and S2, sequentially and repeatedly, under what conditions can anticipatory regulation evolve as an adaptive strategy? To answer this question, we employ two approaches.

In the first, we use a simple mathematical model to represent exposure of a population to two temporal stimuli and identify the network and physiological parameters, which maximize fitness in the given conditions. Based on the inputs from our modelling results, in the second, we evolve *E. coli* under alternating exposure to a pentose sugar rhamnose (S1) and an oxidative stress molecule, paraquat (PQ) (S2).

Briefly, *E. coli* utilizes the transporter RhaT to internalize rhamnose ^20, 21^. Thereafter, the sugar is processed by metabolic enzymes RhaB, RhaA, and RhaD ^22^. The regulon is controlled by two AraC-like transcription factors, RhaR and RhaS ^23–25^. Both bind upstream of the coding sequence of the genes essential for rhamnose utilization. The fact that rhamnose utilization regulon has two regulators and that the coding regions of one of the transcription factors (RhaR) and the transporter (RhaT) overlap make the rhamnose utilization regulon unique in *E. coli*^23, 26, 27^. The rhamnose-utilization regulon is also under catabolite repression, when glucose is present in the environment ^28^.

Prokaryotes protect themselves against oxidative stress (H_2_O_2_, ·OH free radical, ·O^2−^) with the action of superoxide dismutases, catalases, and peroxiredoxin ^29^. In the presence of oxidative stress, the expression of these enzymes is up-regulated by transcription factors, OxyR ^30^, SoxS, and SoxR ^31^. In a recent study, more than 50 distinct binding sites were reported for OxyR, SoxS and SoxR, when *E. coli* was studied under paraquat stress ^32^. Most of these binding sites were found to be RpoD (Sigma 70)-dependent, and combined, the three transcription factors controlled more than 100 genes in response to oxidative stress. The OxyR, SoxRS regulon includes activation of *zwf* (which encodes glucose-6 phosphate-1-dehydrogenase), increase in cellular NADPH pool ^32, 33^, aromatic amino acid production, and upregulation in cell wall biosynthesis ^34^.

Repeated, alternating exposure to S1 and S2 for ∼850 generations, leads to evolution of anticipatory regulation, where prior exposure to rhamnose provides an adaptive benefit when the population is shifted to paraquat. This benefit is observed only in lines which were exposed to alternating S1 and S2; and is only seen when the cells are pre-exposed to rhamnose. Gene expression studies demonstrate that one of the possible mechanisms of this adaptive benefit is partial activation of *soxS*, upon exposure to rhamnose. Our study thus demonstrates that, in controlled laboratory environment, anticipatory gene regulation can evolve in a short timeframe of a few hundred generations.

## Methods

### Model system description

Two regulatory modules and their possible interaction was studied as follows. A microorganism experiences an environmental signal S1. Upon sensing this signal, it modulates gene expression. As part of this response, a transcription factor R1 gets activated and triggers expression of a target protein T1. The protein T1 is assumed to serve a physiological function, which helps the bacterium survive in S1. The signal S1 is assumed to be present for time *t1*. After this time, S1 is replaced by another signal S2. In response to S2, the cell activates a transcription factor R2. The activated R2 then positively controls expression of the target protein T2, which helps the cell survive and replicate in S2. The signal S2 persists for time *t2* (**Figure 1**).

**Figure 1.**
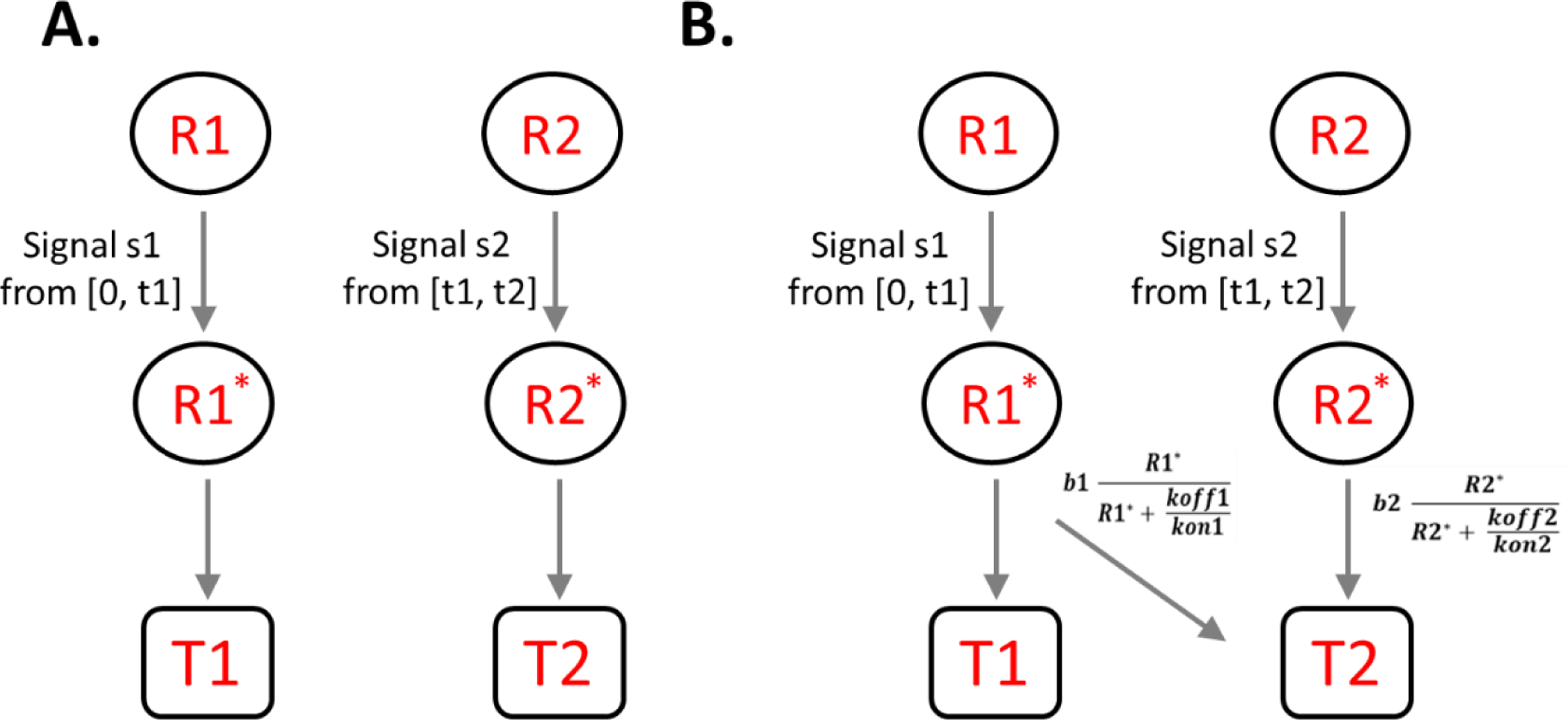
Regulatory designs without (A) and with (B) anticipatory regulation. In the prevailing environment, signal s1 is present from [0, t1] and signal s2 is present from [t1, t2]. In the first design, activated regulator R1* controls expression of target protein T1, and R2* controls expression of T2. In the second regulatory design, R1*, in addition to controlling expression of T1, also partially/fully controls expression of target protein T2.

In the first regulatory design, it is assumed that there is no regulatory crosstalk between the two signal responses. That is, activated R1 is responsible for response to S1 (controlling expression of T1), and activated R2 is responsible for response to S2 (controlling expression of T2). We call this design a no-anticipatory regulation design (Figure 1A). In such a setting, when allowed to propagate in alternating S1 and S2 for a long time, the population will evolve and enhance fitness. In our mathematical framework, we solve for parameter values, which maximize fitness for the regulatory topology in Figure 1A.

On the other hand, in the design with anticipatory regulation, the promoter of T2 is controlled by active R1 as well as active R2 (Figure 1B). We solve for the values of these biochemical interaction that maximize fitness in this regulatory design.

In particular, we are interested in identification of parameter values such that the fitness conferred by design in Figure 1B (anticipatory regulation) exceeds that by design in Figure 1A (no anticipatory regulation).

The equations dictating the synthesis and degradation rate of proteins in the cell, upon sensing signal S1 or S2 can be written as below.

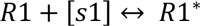

and,

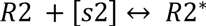

is represented as:

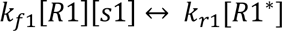

And

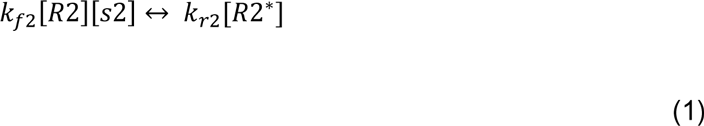

The dynamics of target proteins T1 and T2 is represented as the following equations. *R1\** is the transcription factor responsible for expression of T1, *Km* is the Michaelis-Menten kinetic constant for T1 production, *b_o_* is the maximum production rate of T1, and *k_d_* is the combined degradation and dilution rate of T1 in the cell ^35^. In our analysis, we assume that expression of R1 is not contingent on the presence or absence of signal S1 (but its activation is).

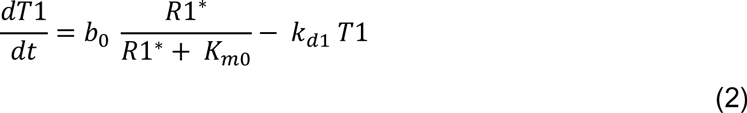

Similarly, for design in Figure 1A, expression of T2 is under the exclusive control of R2.

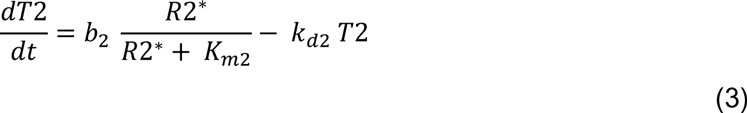

However, in the case when anticipatory regulation controls gene expression (Figure 1B), the dynamics of T2 in the cell can be quantified as:

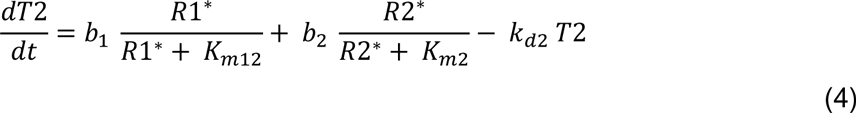

Where,

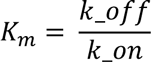

for each transcription factor-DNA interaction. In this manner, given the biochemical parameters of a network, we can quantify the dynamics of protein expression in a cell.

### Cost-benefit framework

From the time-course data of protein expression, we use a cost-benefit model to understand the cellular fitness, as a result of the gene expression dynamics ^36^. Such a framework has been used previously to understand cellular behavior and physiology ^37–39^. In this formulation, the benefit conferred to the cell by the target protein T2 is given by the following expression:

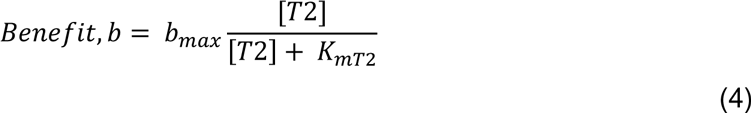

In addition, protein production is known to have an energetic cost associated with it ^40^. This cost is a linear function of the amount of protein being synthesized. Hence, the cost component associated with synthesis of protein T2 can be represented as:

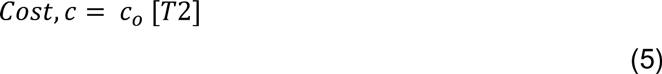

The overall fitness of a cell is simply the difference of the benefit and cost associated with synthesis of T2. Hence, fitness is given by:

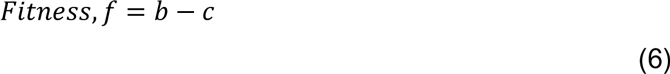

Given a set of network topology and the biochemical parameters associated with it, this framework allows us to compute the benefit of production of T2 and track this quantity with time. In the context of this work, we compute the fitness conferred by T2 in the time window [*t1, t2*] and the total cost associated with synthesis of protein T2 in the window [*t0*, *t2*]. Note that synthesis of T2 in the regulatory design with anticipatory regulation starts before time *t1*.

### Solving for optimal fitness

Equations (1-6) have biochemical constants appear as equation parameters. Each parameter represents specific cellular processes. For instance, *Km* represents the interaction dynamics between the transcription factor and DNA. The parameter *b* represents the maximal rate of transcription from the promoter of gene *t2*.

In this work, we assume that the parameters associated with changing the regulatory patterns of gene expression (like, *Km* and *b*) are evolvable on a time scale much faster than the time needed to change parameters associated with protein property (like, enzyme activity, degradation rate) ^41^. This has been shown in several studies ^42–44^. As a result, we let the parameters, which control regulation evolve, while keeping the others constant. Moreover, the parameters associated with these biochemical functions cannot take arbitrary values. Each of these parameters is constrained by the thermodynamics associated with biological processes. Hence, we let the parameters change within a defined range (given in **Table 1**). The range chosen for each parameter represents the biologically permissible window in which they can take a value ^45–49^.

**Table 1.** Range of values of the parameters used in this study.

| Parameter | Value/Range |
| --- | --- |
| $b$ (for all promoters) | 0 to 0.3 |
| $k_{on}$ (for all DNA-protein interactions) | 0 to 60 |
| $k_{off}$ (for all DNA-protein interactions) | 0.3 |
| $kd$ (degradation rate constant) | 0 to 0.015 |
| Signal (S1 or S2) | 0 or 1 |
| $b_{max}$ (maximum benefit) | 1.5 |
| $Km$ (half-maximum benefit) | 15 |
| $co$ (cost of synthesis of one protein molecule) | 0.01 |

Given the above, we introduce “mutations” (change parameters) in the network and for each of the two regulatory designs (with and without anticipatory regulation, as defined above), find the parameter set which optimizes fitness. We are particularly interested in finding whether fitness is optimized via topology in Figure 1A or 1B. Note that in this scheme, when anticipatory regulation is not the optimal response, *b_12_* = 0 should be optimal design in the regulatory topology.

All simulations were performed in Matlab 7.0.

### Strain used

*E. coli* K12 MG1655 (ATCC 47076) (F-lambda-) was used in this study. The *soxS* mutant was generated using the primers 5’-CCCCAACAGATGAATTAACGAACTGAACACTGAAA AGAGG GTGTAGGCTGGAGCTGCTTC-3’ and 5’-GAGCAATTACCCGCGCGGGAGTTAACGCGCGGGCAATAAACATATGAATATCCTCCTTA-3’ from the parent strain, as per the method described by Datsenko and Wanner ^50^.

### Media and reagents

M9 glycerol medium contained 0.2% glycerol unless otherwise stated. M9 salts, trace elements and casamino acids were prepared in concentrated stocks. Stock of a 5x M9 salts solution consisted of 64 g/L Na_2_HPO_4_.7H_2_O, 15 g/L KH_2_PO_4_, 2.5 g/L NaCl, and 5 g/L NH_4_Cl dissolved in Milli-Q filtered water. Casamino acids were prepared as 10x solution and were used at a final concentration of 0.05% (w/v) in the growth media. MgSO_4_ and CaCl_2_ were prepared at 1M stock solutions each. All stock solutions were sterilized by autoclaving. The stocks of rhamnose and paraquat was prepared at 20% and 5mM respectively and sterilized by filtration through 0.22 μm filters.

### Laboratory evolution experiment

Adaptive evolution experiments were performed by serial dilutions every 12 hours in M9 medium with appropriate stimulus. All cultures were grown at 37°C and 250rpm unless otherwise stated. Every 12 hours, growing cultures of bacteria were diluted 1:100 in fresh M9 medium with appropriate stimulus yielding ∼6-7 generations every 12 hours. The choice of sub-cultures every 12 hours ensured that cells were transferred to fresh media before they entered stationary phase. This was done to avoid the physiological changes that the bacteria undergo, upon entering the stationary phase ^51^. Moreover, RpoS is also known to confer stress resistance ^52^. The evolution experiment was carried out for a total of 850 generations. Freezer stocks of intermediate time points were made in the experiment. Analysis of the stocks at 300 and 600 generations are presented in this work. Starting from the ancestor, three independent lines were evolved for each environmental condition.

### Conditions for evolution of anticipatory regulation

We used rhamnose and paraquat at a concentration of 0.2% and 40 μM respectively, as the two stimuli. The evolution experiment was carried out in three replicate lines serially diluted 1:100 in alternating conditions of rhamnose and paraquat every 12 hours, as described above. As controls, we evolved three independent lines in either 0.2% rhamnose only or 40uM paraquat only. The control experiment were both the stimuli were added together would not grow beyond four dilutions, hence was dropped out of the experiment.

### Analysis of evolved lines for anticipatory regulation

All evolved lines and the ancestor were revived from freezer stocks into 2ml LB and incubated for 12 hours with shaking at 250 rpm at 37°C. The cultures were then sub-cultured 1:100 in 2ml M9 medium, containing 0.2% glycerol as the carbon source, for 12 hours. From each tube, cells were then sub-cultured 1:100 in 2ml M9 glycerol media (a) with and (b) without 0.2% rhamnose, and allowed to grow for 12 hours at 250 rpm at 37°C. After growth for 12 hours, all lines were sub-cultured 1:100 into 2ml M9 with PQ (40uM), containing glycerol as the carbon source. A volume of 150 μL of these cultures were transferred to a 96-well clear flat-bottom microplate (Costar) in triplicates. The cultures were grown at 37°C in an automatic microplate reader (Tecan Infinite M200 Pro), until they reached stationary phase. OD600 readings were taken every 30 minutes with 10 minutes of orbital shaking at 5mm amplitude before the readings. A gas permeable *Breathe-Easy* (Sigma-Aldrich) sealing membrane was used to seal the 96-well plates.

### Gene expression measurements

The 850 generation lines from each lineage and wild type were transformed with *soxS*-GFP, a plasmid-based promoter fusion from Thermo Scientific *E. coli* promoter collection (PEC3877) ^53^. For estimating the promoter activity under different conditions all evolved lines with *soxS*-GFP plasmid and the wild-type ancestor were revived from freezer stocks from the −80°C deep freezer into 2ml LB and incubated for 12 hours at 37°C. All the media for these transformed lines contained 40µg/ml Kanamycin. The saturated cultures were then sub-cultured 1:100 in 2ml M9 glycerol medium for 12 hours. All lines were then sub-cultured 1:100 into fresh M9 glycerol medium without/with 0.2% rhamnose and allowed to grow for 12 hours at 37 deg C. After this period of growth, 150 μL of these cultures was transferred to a 96-well Black clear flat-bottom microplate (Costar) and Fluorescence (488/525 nm) and OD600 readings taken using a microplate reader (Tecan Infinite M200 PRO).

For *soxS*-GFP expression in paraquat, cells were transferred at a 1:100 dilution from M9 glycerol to M9 glycerol with /without 0.2% rhamnose. The culture were then allowed to grow for 12 hours at 37°C. The resulting cultures were diluted to an initial OD600 of 0.1 in 2ml M9 glycero media with 40μM PQ. A volume of 150 μL of these cultures were transferred to a 96-well Black clear flat-bottom microplate (Costar) in triplicates. The cultures were grown at 37°C in an automatic microplate reader (Tecan Infinite M200 Pro), until they reached stationary phase. Fluorescence (488/525 nm) and OD600 readings were taken every 30 minutes with 10 minutes of orbital shaking at 5mm amplitude before the readings. A gas permeable *Breathe-Easy* (Sigma-Aldrich) sealing membrane was used to seal the 96-well plates.

### Sequencing

Promoters of four target genes of OxyR and SoxSR were sequenced. The primers used for sequencing were as follows: *zwf* (CCAGCATATTCATGATGTAA and GTCATTCTCCTTAAG TTAAC); *sodA* (TAATCGCGTTACTCATCTTC and ATTCATCTCCAGTATTGTCG); *fpr* (CTGATTGATTTGATCGATTG and GTTTTTCTCCTGTTTTGATT); and *fumC* (TGAAAT AAACAGAGCCGCCC and GACCTGCTCCTCACCTGATT). For sequencing, individual colonies of respective strains were collected from LB plates, and allowed to grow for 4-6 hours in liquid LB media at 37 deg C. The cells were harvested, and their genomic DNA isolated. The four promoter sequences were amplified by PCR. Sequencing was done by Eurofins Scientific.

## Results

### A framework to identify when anticipatory regulation can evolve in transcription networks

We simulate a scenario where a bacterium is responding to two environmental cues encountered in temporal succession. The response to the first cue, S1, is expression of target protein T1 and the response to the second cue, S2, is expression of target protein T2 (Figure 1A). As a starting point, we assume that expression of T1 and T2 (in presence of the respective cues) is triggered by transcription factors R1 and R2, respectively. Under what conditions, however, is this regulatory design rewired such that transcription factor R1, in addition to activating expression of T1, also controls expression of T2 (Figure 1B)? In other words, under what conditions associated with the environments or properties of the proteins involved does the design represented by Figure 1B confer a greater fitness as compared to one in Figure 1A?

### Anticipatory regulation is an adaptive response for intermediate values of degradation rates of the protein T2

We simulate the networks in Figure 1A and 1B for a set of parameter values, chosen from a window typically representing each biochemical interaction in a cell ^45–49, 54, 55^. We focus on control of transcription as the most flexible parameters of genetic rewiring of networks, as promoter mutations are thought to be accomplished much quicker than protein property changes (like, enzyme activity) ^41^. For a given value of *kd* (degradation + dilution rate constant for protein T2), we find the regulatory design in Figure 1, which leads to a maximal fitness.

For a given set of fixed parameters, maximal fitness is obtained via three distinct regulatory logics, depending on the degradation rate of the protein T2 (**Figure 2A**). For small values of the *kd* of T2, the maximal fitness is achieved when control of T2 is entirely under the regulator R1. That is, R2 does not play any role in controlling expression of T2. This can be understood as follows. Expression of T2 by R1 leads to T2 production even before signal S2 is present in the environment. As a result, the T2 so produced does not confer any advantage to the cell. Thus, this preemptive production of T2 does not give any benefit to the cell, but does come at an additional cost associated with protein production in the cell. This cost, however, is offset by the enhanced T2 levels inside the cell (and consequently the benefit conferred to the cell), from the time when signal B is introduced to the system. When T2 is stable, the steady state T2 in the cell is high, and as a result, the fitness of the cell at the moment signal B is introduced to the system is also high.

**Figure 2.**
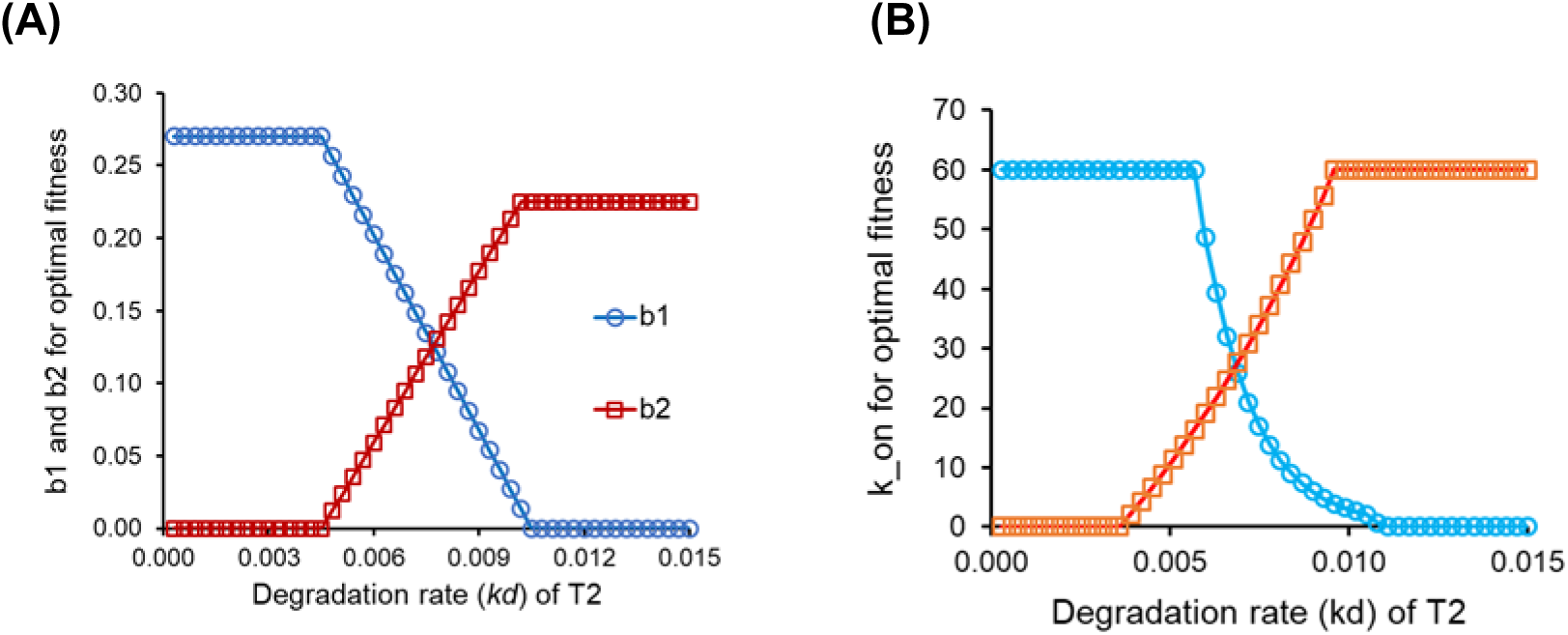
**(A) For intermediate values of the degradation rate *kD* for target protein, T2, conditioning is the optimal response.** For low values of the degradation constant, expression of T2 controlled entirely by T1 (i.e. b2 = 0) is the optimal response. For high degradation rates, optimal fitness is obtained when expression of T2 is controlled by R2 (i.e. b1 = 0). [b1 is the maximal promoter strength when R1* triggers expression from promoter of T2; b2 is the maximal promoter strength when R2* triggers expression from promoter of T2]. **(B) For intermediate values of the degradation rate *kD* for target protein, T2, conditioning is the optimal response.** For low values of the degradation constant, expression of T2 controlled entirely by T1 is the optimal response. For high degradation rates, optimal fitness is obtained when expression of T2 is controlled by R2 (i.e. k_on for b1 = 0). Blue circles represent the k_on for interaction between R1* and promoter of T2. Orange squares represent k_on for interaction between R2* and promoter of T2.

At the other extreme, when the *kd* of T2 is high, the regulatory design that optimizes cellular fitness is one where control of expression of T2 is not by R1, but is instead entirely by R2. In such a context, the energetic cost of T2 production in the “futile” time-period [0, *t1*] prior to introduction of signal B, because of the high protein turnover, is high. In such a scenario, the cellular adaptive strategy is control of each target protein by its cognate regulatory partner.

However, at intermediate values of the protein T2 degradation rate constant, for maximal cellular fitness, both b1 and b2 are non-zero. This window of the values of the degradation rate constant of T2 represents the scenario where conditioning is the adaptive response in cellular functioning. Such a set up identifies conditions for a distributed regulatory design for control of gene expression. Interestingly, distributed control strategies are known to be more robust in control systems, and have also been shown to be present in biological regulatory designs ^56^.

Among the three regimes, anticipatory regulation is only observed when b1 and b2 are both non-zero. The other two regimes are solutions to the optimization problem, which lead to different regulatory designs. Thus, one of the key variables which leads to anticipatory regulation as the optimal solution is the degradation rate of the target protein.

In the above result, we only compare the optimal solutions of the two regulatory designs. That is, we locate the parameter values of the two regulatory designs (with and without anticipatory regulation), which optimize fitness. In such a context, we note that anticipatory regulatory design can outperform the other under certain parameter regimes. Accessibility of these optimal solutions via an adaptive path on a landscape, however, is not known. Walks on adaptive landscapes is a multidimensional problem and little is known about structures of an actual fitness landscape ^57^. In the context of the problem we discuss, how a mutation changes the parameter values is an open problem ^58^. Therefore, from a starting parameter set, to what adaptive solution does a population evolve to is an open problem, and has been observed to depend on the details of the problem. Little is known about this facet of evolutionary dynamics ^59^.

Another scheme to incorporate changes in regulatory design would be to not change the maximum strength of the promoter, but change the binding affinity of the transcription factor with the promoter region of the DNA. In the context of the model, we change the parameter k_on, which represents the affinity of the DNA for the transcription factor.

Similar to the results obtained above, we note that anticipatory regulation could also outperform for intermediate values of the degradation rate of protein T2, when we permit k_on1 or k_on2 to vary (**Figure 2B**). Also similar with the previous results, while conditioning only outperforms the non-conditioning designs for intermediate values of *kd* of protein T2, positive fitness in regulatory designs with conditioning exist for all values of kd.

We demonstrate that anticipatory regulation offers the optimal fitness in certain scenarios, for the simplest gene regulatory circuit. It is likely that as we move towards regulatory topologies of greater complexities, the ability to incorporate newer features in the dynamical responses of networks also increases. These responses could include cell-cell heterogeneity, memory, speed of response among others. All these properties have shown to have fitness consequences for a cell, in different contexts ^39, 55, 60, 61^. In addition, the number of ways in which these responses could be incorporated into the cellular manifestations also increases as the size of the network increases. This facet of networks makes tracking evolutionary dynamics of a system quite daunting – not only in terms of <u>how</u> a design could incorporate a specific feature into cellular response, but also in terms of what additional features (and also future possibilities) are going to get incorporated in the cellular physiology as a fallout of the adaptive response.

### Anticipatory regulation, though not optimal, confers a positive fitness to the cell

Figure 2 shows that anticipatory regulation is not the optimal response for very low and very high values of the degradation and dilution rates for protein T2. This is further illustrated in **Figure 3**. Here, ΔF corresponds to the difference between the best fitness when anticipatory regulation is permitted (for a corresponding (b1, b2) or (kon1, kon2) pair) and the best value of fitness when it is not permitted. Thus, for a high degradation and dilution rate (which lies in the region where regulatory design in Figure 1A maximizes fitness), ΔF < 0 for all combinations of (b12, b2) (**Figure 3A**) and (k_on12, k_on2) (**Figure 3B**).

**Figure 3.**
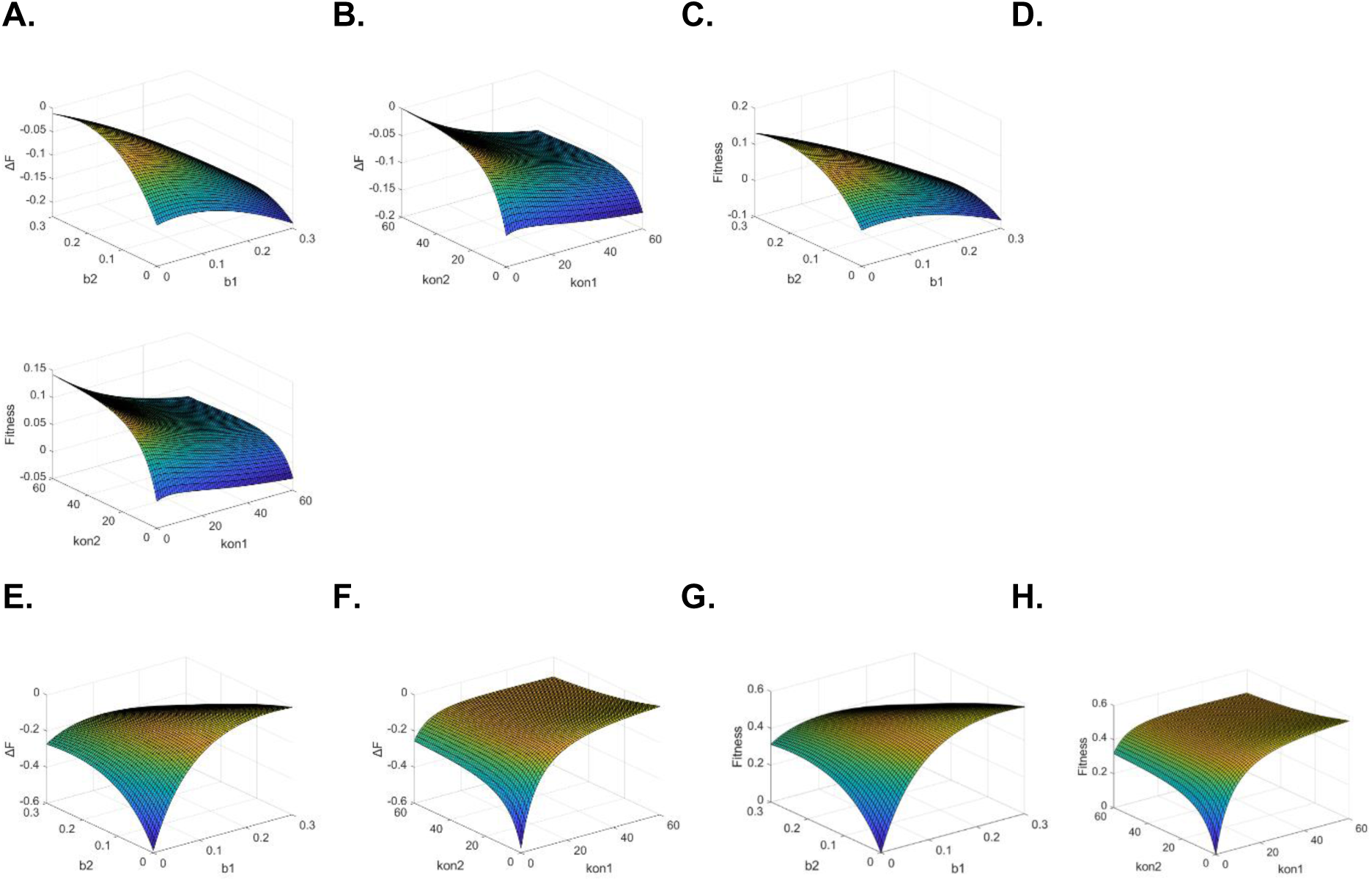
**Anticipatory regulation yields positive fitness at high protein degradation and dilution rates (A-D).** When the promoter strength **(A)** or the DNA-protein interaction **(B)** is allowed to change, the fitness of anticipatory regulatory logic is always less than of a non-anticipatory regulatory design in Figure 1A. However, for large parts of the range of parameter values, the absolute fitness conferred in the regulatory design corresponding to conditioning is greater than zero **(C-D)**. **Anticipatory regulation yields positive fitness at low protein degradation rates (E-H).** When the promoter strength **(E)** or the DNA-protein interaction **(F)** is allowed to change, the fitness of conditioning regulatory logic is always less than the design when R1* controls both T1 and T2 gene expression. However, for large parts of the range of parameter values, the absolute fitness conferred in the regulatory design corresponding to conditioning is greater than zero **(G-H)**. (A-D) are for *kd* (for T2) equal to 0.0144. (E-H) are for *kd* (for T2) equal to 0.0025.

However, for a large parts of the parameter scanned in these heat plots, the fitness when anticipatory regulation is permitted, is greater than zero (Figures 4C and 4D). Hence, even though, it is not the best possible response in these parameter regions, it may still be observed as the prevailing regulatory design. This is particularly so when strength of selection on the target protein is not very high, or in population regimes where drift has a significant role to play.

**Figure 4.**
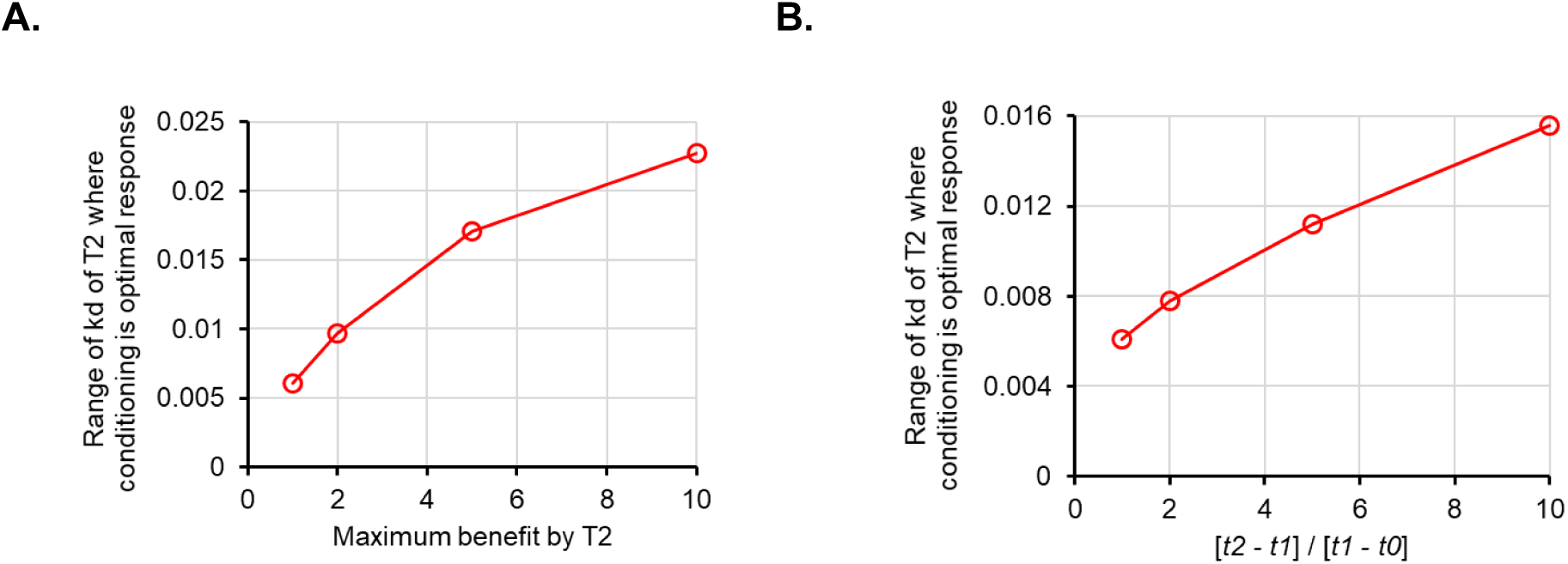
Anticipatory regulation emerges as optimal solution as (A) Maximum benefit conferred by T2 increases, and (B) Relative time of S2 with respect to S1 increases. The y-axis on the plots represents the range of the values of *kd* for protein T2, for which anticipatory regulation is the optimal regulatory design.

Similar results are also observed when parallel heat plots are analyzed for protein-DNA interaction of T2. We note that while anticipatory regulation might not be the best possible response, it corresponds to greater than zero fitness values of the system, for a large part of the parameter regime (**Figure 3E-H**).

### Can anticipatory regulation be evolved in a laboratory set-up?

Given the above, we speculate the requirements of an experimental design to demonstrate evolution of anticipatory regulation in laboratory conditions. Our model lays down conditions, which might facilitate design of experiments towards this goal.

We predict that anticipatory regulation is most likely to evolve if the target protein has an intermediate degradation and dilution rate. This is perhaps the single most important criteria for anticipatory regulation to emerge as an outcome of an evolutionary experiment. In addition to this, the target protein responding to condition/environment 2 should serve a physiological function, which enhances cellular fitness considerably. Alternatively, the absence of target protein T2 should cause a significant fitness cost to the cell (**Figure 4A**). In this context, for experimental evolution of anticipatory regulation, it makes sense if the design of experiment is such that the signal S2 is a stress, which, in the absence of the appropriate cellular response, has a significant fitness penalty to the cell.

The other variable which dictates the likelihood of evolution of anticipatory regulation is the relative timing of the two environmental signals. In this context, we note that the greater the relative time for which the signal S2 prevails in the environment, the greater the likelihood of conditioning evolving as the optimal fitness solution (**Figure 4B**). Thus, in the context of design of experimental evolution, the relative timing of the two signals is another key variable, which likely dictates the evolutionary fate of the population. From the context of bacteria in their ecological niches, cyclical nature of the environment is not too stringent a requirement to be fulfilled for anticipatory regulation to emerge. Organisms in ecological niches are exposed to cyclical daily environments, and hence, selection oscillates with time.

However, these considerations are unlikely to be sufficient for evolution of anticipatory regulation. Moreover, while we only analyze incorporation of anticipatory regulation via transcription, several other possibilities remain. Gene regulation can be incorporated at via post-transcription, translation, and/or post-translation mechanisms. These regulatory possibilities only extend the number of ways that anticipatory regulation can be built into a regulatory design of cellular networks, should it confer adaptive benefit to the cell.

Timescale of evolution of anticipatory regulation also remains unknown. It depends on the selection pressure we put, the choice of S1 and S2, and the background genotype on which the experiment is performed.

### Evolution of anticipatory regulation in *E. coli*

From our modeling exercise, we note that two key parameters for evolution of conditioning are (a) an intermediate degradation and dilution rate of the target T2, and (b) a large cost associated with not initiating a rapid response to signal S2. In view of these inputs, we chose a pair of environmental signals, which satisfy these conditions. We chose the pentose sugar rhamnose as signal S1 and an oxidative stress molecule, paraquat (PQ) as signal S2. Rhamnose utilization is controlled by two transcription factors, RhaS and RhaR^62, 63^. In the presence of rhamnose, RhaR activates the expression from the *rhaSR* promoter^64^. RhaS, on the other hand, activates expression of the rhamnose transporter protein, RhaT, and the catabolic operon, *rhaBAD*^27^. Both, RhaR and RhaS, belong to the AraC/XylS family of transcriptional regulators, and share sequence similarity in the DNA binding regions of the proteins^65–67^.

On the other hand, oxidative stress response is largely controlled by OxyR and SoxRS system in *E. coli*^68^. To the best of our knowledge, the two systems are not linked by any direct transcriptional regulation. A stress of 40 uM PQ induces a growth defect in *E. coli* (**Figure 5A**). Thus, any anticipated expression of the SoxRS regulon is likely to provide a large adaptive benefit to the individual. With this premise, we performed an evolutionary experiment where the cells were exposed to S1 and S2 alternatively for a total of about 850 generations (See experimental design in **Figure 5B**, and methods section). Alongside, we also performed experiments where populations were exposed to only S1 or only S2.

**Figure 5.**
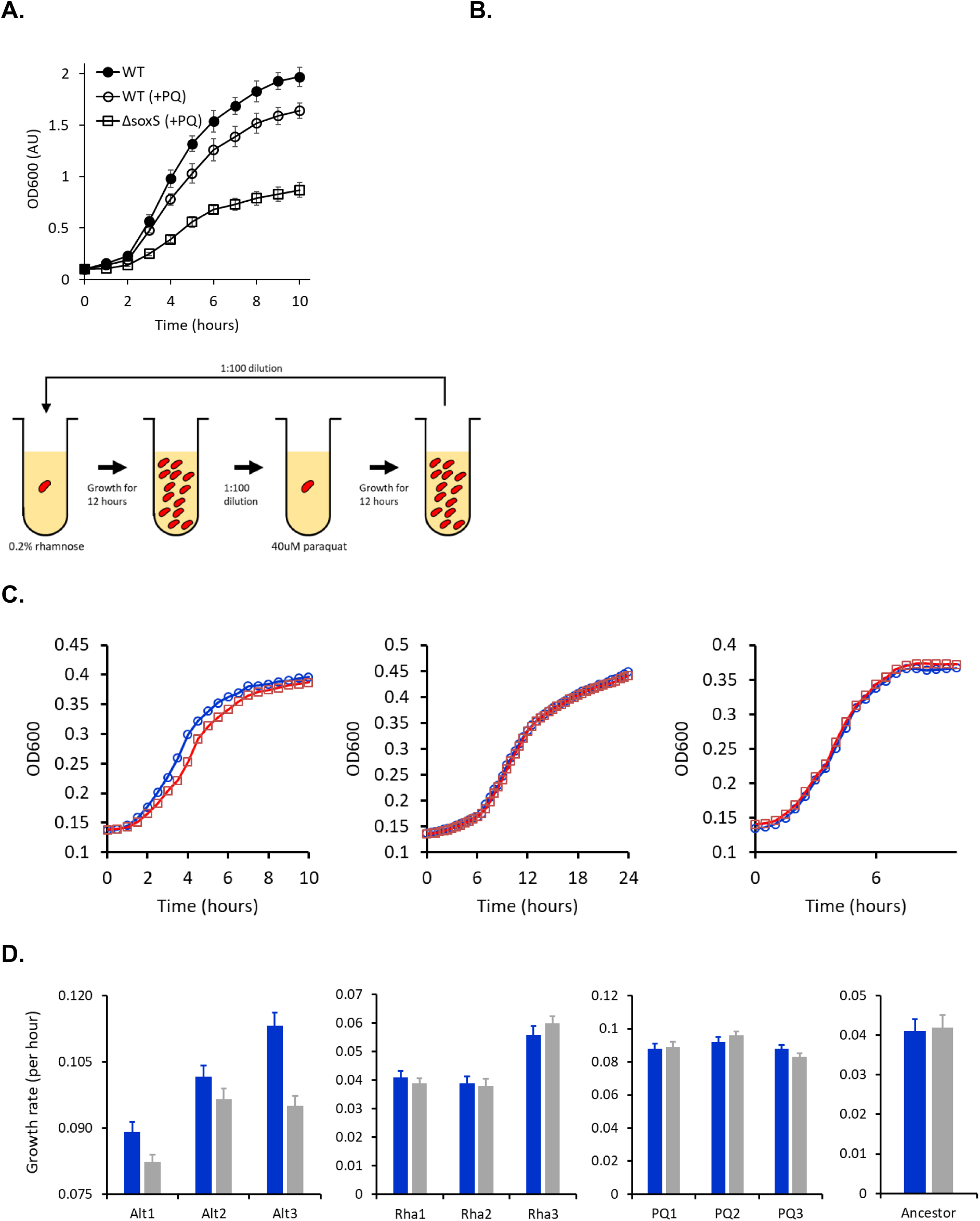
**(A)** *E. coli* exhibits a growth defect in M9 glucerol media, when grown in the presence of 40 μM PQ. **(B)** Experimental plan for the rhamnose-PQ alternating evolution. Cells were exposed to rhamnose and then PQ for 6-7 generations for a total of 850 generations. **(C) Anticipatory regulation evolves in rhamnose-PQ alternating lines.** Cells from rhamnose-PQ evolved line (Line Alt1) (left), rhamnose-only evolved line (Line Rha1) (middle), and PQ-only evolved line (PQ1) (right) were grown in M9 minimal media with (blue) or without (red) rhamnose. The cells were transferred 1:100 to M9 glycerol media with 40 μM PQ, and their growth kinetics observed. **(D) Anticipatory regulation provides a growth advantage in all rhamnose-PQ alternating evolved lines.** Growth rates of the evolved lines in M9 glycerol media containing 40 μM PQ, when transferred from M9 glycerol media with (blue) or without (grey) 0.2% rhamnose.

In such a context, conditioning can be said to have evolved when, after alternating exposure to rhamnose and PQ for ∼850 generations, pre-exposure to rhamnose confers a growth advantage when the cells are exposed to PQ. Moreover, this advantage should only be present in the lines, which were evolved in an environment with alternating presence of rhamnose and PQ. This advantage should be absent in the lines which were exposed to only rhamnose, and the lines which were exposed to only PQ.

In **Figure 5C**, growth kinetics of one of the lines from the three experimental conditions (Lines 1 from alternating rhamnose-PQ; only rhamnose; and only PQ) are shown. When cells evolved in alternating rhamnose-PQ were pre-grown in rhamnose, they exhibit a growth advantage. This advantage is absent in the lines evolved in rhamnose-only, or PQ-only.

In **Figure 5D**, the growth rate for all three lines in each of the three environments is shown. Among all the lines, only rhamnose-PQ alternating evolved lines show a growth advantage, when pre-exposed to rhamnose. The relative amounts of the advantage vary between the three lines. As expected, the rhamnose-evolved lines show a considerably lower growth rate as compared to the PQ-evolved and the rhamnose-PQ alternating evolved lines.

At 850 generations, the rhamnose-PQ alternating evolved lines exhibit a growth advantage due to anticipatory regulation. To test the kinetics of the growth advantage phenotype in the course of our evolution experiment, two intermediate time-points (generation 300 and 600) in the evolutionary experiment were also checked for their growth rate. This was done as described in Figure 5C. As shown in **Figure 6**, the anticipatory regulation was not observed at the 300 generation time point. By 600 generations, anticipatory regulation was observed in two of the three rhamnose-PQ alternating lines. This trend was exaggerated by generations 850 (**Figure 6**). The growth advantage, associated with a prior exposure to rhamnose, is at both the intermediate points, is not observed in rhamnose-only and PQ-only evolved lines. Interestingly, in two of the three rhamnose-evolved lines, fitness decreases in the PQ environment. Trade-off between performance in two different environments has been studied extensively in recent years ^69–71^.

**Figure 6.**
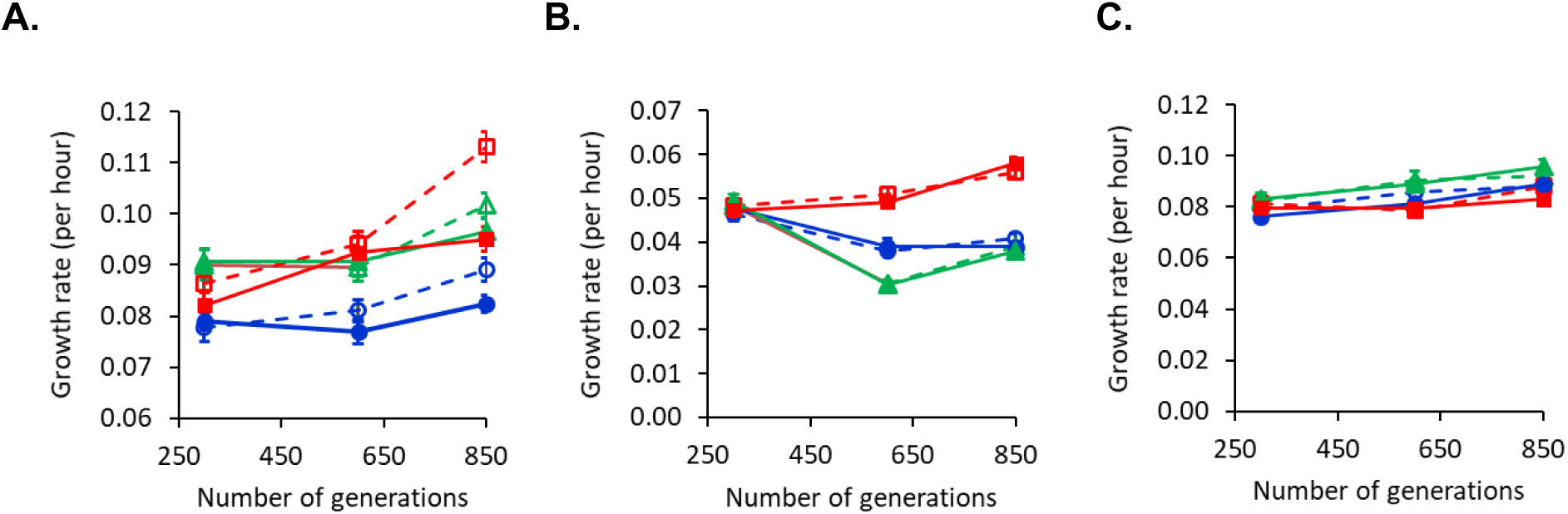
Growth advantage because of evolution of anticipatory regulation takes place around and after 600 generations into the experiment. Growth rate of Lines 1 (Blue), 2 (Green), and 3 (Red) in rhamnose-PQ alternating conditions (**A**), rhamnose-only (**B**), and PQ only (**C**). Solid and dashed lines represent growth curve of the lines, when transferred from M9 glycerol medium without or with rhamnose, respectively.

### Molecular targets leading to evolution of anticipatory regulation

OxyR and SoxRS are the two major transcription regulatory systems, responsible for cellular response to oxidative stress. The combined regulon of these two systems is more than a 100 genes in *E. coli.* For identification of regions of DNA where a mutation could have facilitated evolution of anticipatory regulation, we sequenced the promoter regions of four target genes which respond to oxidative stress. These genes include *zwf* (enzyme in the pentose phosphate pathway and a response to oxidative stress) ^72–75^, *sodA* (one of three superoxide dismutases in *E. coli*) ^76–78^, *fumC* (one of the three fumarase isozymes participating in TCA and is expressed in response to oxidative stress) ^79–81^, and *fpr* (Flavodoxin/ferredoxin-NADP+ reductase) ^82, 83^ promoters. The promoter regions of these four genes were sequenced in the three lines evolved in rhamnose only, PQ only, and rhamnose-PQ alternating lines.

In these sequenced parts of the genome, two mutations were identified when compared with the ancestor’s genotype. Both these mutations were found in the lines evolving in rhamnose-PQ alternating conditions. In one of the lines (Alt1), in the region upstream of the coding sequence of *zwf* an adenine residue 57 bases downstream of the transcription start site of the gene was deleted. This nucleotide likely interferes with the mRNA structure and stability. The mutation could also alter the ribosome binding site of the *zwf* mRNA, and hence change the translational efficiency of the gene. In another line evolved in the rhamnose-PQ alternating environment (Alt2), an insertion (adenine) was identified 46 bases downstream of the transcriptional start site of the *zwf* gene. As a result of this mutation, the inhibitory function of small RNAs, fnrS and rhyB, is likely compromised ^84, 85^. The rhamnose-only and PQ-only evolved lines did not have any mutations in the promoters sequenced in this work. Evolution of anticipatory regulation likely involves a large set of mutations. Compared to the ancestral growth rate in PQ (Figure 5D), the growth rate of the evolved lines by generation 600 is considerably higher. However, this increase in the growth rate is achieved without anticipatory regulation. Thus, our data suggests that a set of mutations are likely to have been acquired in the first phase of the evolution experiment. These mutations confer benefit by altering gene expression in a manner independent of anticipatory regulation. Only when these beneficial mutations are fixed in the population, does anticipatory regulation evolve in the culture.

### Evolved lines lead to faster induction of SoxS

SoxS is one of the primary proteins responsible for cellular response towards oxidative stress ^86^. To test the possibility if induction of SoxS differs in the evolved and ancestral lines, when the cells are exposed to rhamnose, we performed the following experiment. *E. coli* is grown in M9 glycerol media in the absence and presence of 0.2% rhamnose. As shown in **Figure 7**, in all three lines evolved with alternating exposure to rhamnose and paraquat, *soxS* promoter is activated in presence of 0.2% rhamnose. This rhamnose-dependent activation of the *soxS* promoter is absent in the lines evolved in rhamnose only, or the lines evolved in paraquat only. In addition, rhamnose-dependent activation of the *soxS* promoter is absent in the ancestor strain.

**Figure 7.**
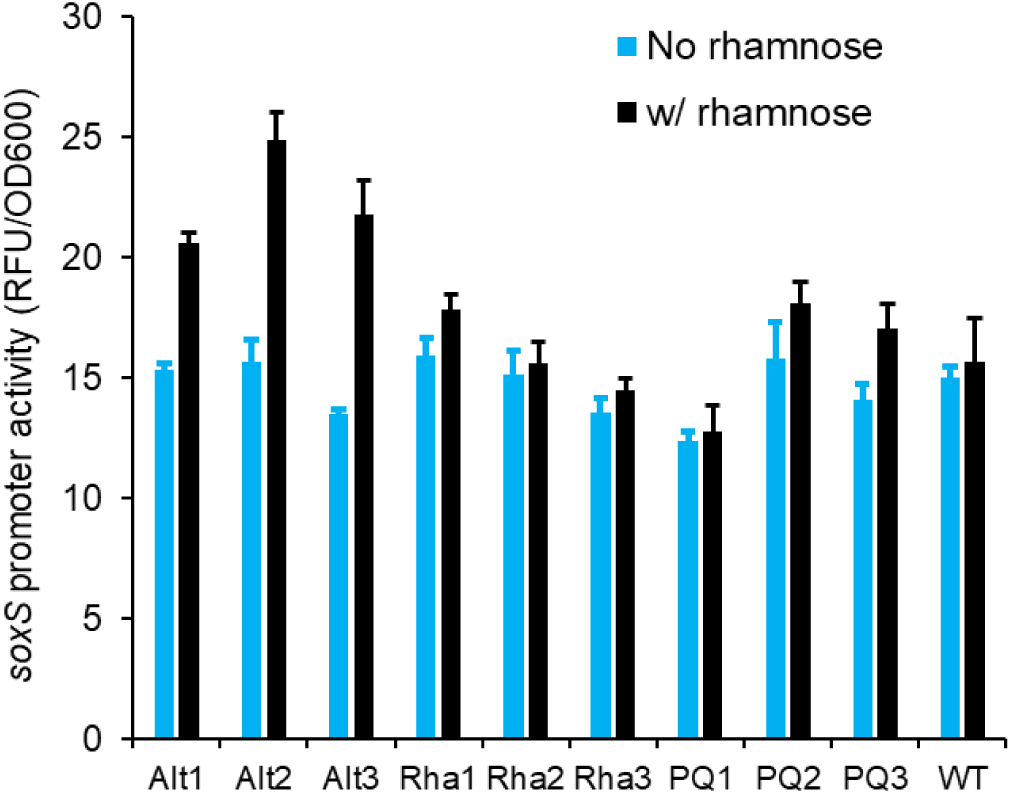
Lines evolved with alternating exposure to rhamnose and paraquat Alt1, Alt2, and Alt3) exhibit a statistically higher expression from the *soxS* promoter in the presence of rhamnose (P<0.005, t-test). This rhamnose-dependent activation of the *soxS* promoter is absent in the ancestor (WT), in rhamnose-evolved lines (Rha1, Rha2, and Rha3), and in paraquat-evolved lines (PQ1, PQ2, and PQ3).

Next, lines Alt1, Alt2, and Alt3 were grown in M9 glycerol media with and without rhamnose. Upon saturation, the cells were transferred to M9 glycerol media containing paraquat. As shown in **Figure 8**, the cells transferred from environments with rhamnose exhibited a faster induction of the *soxS* promoter. This phenomenon was absent in the lines evolved in cyanate, in lines evolved in rhamnose, and in the ancestor (Figure 8).

**Figure 8.**
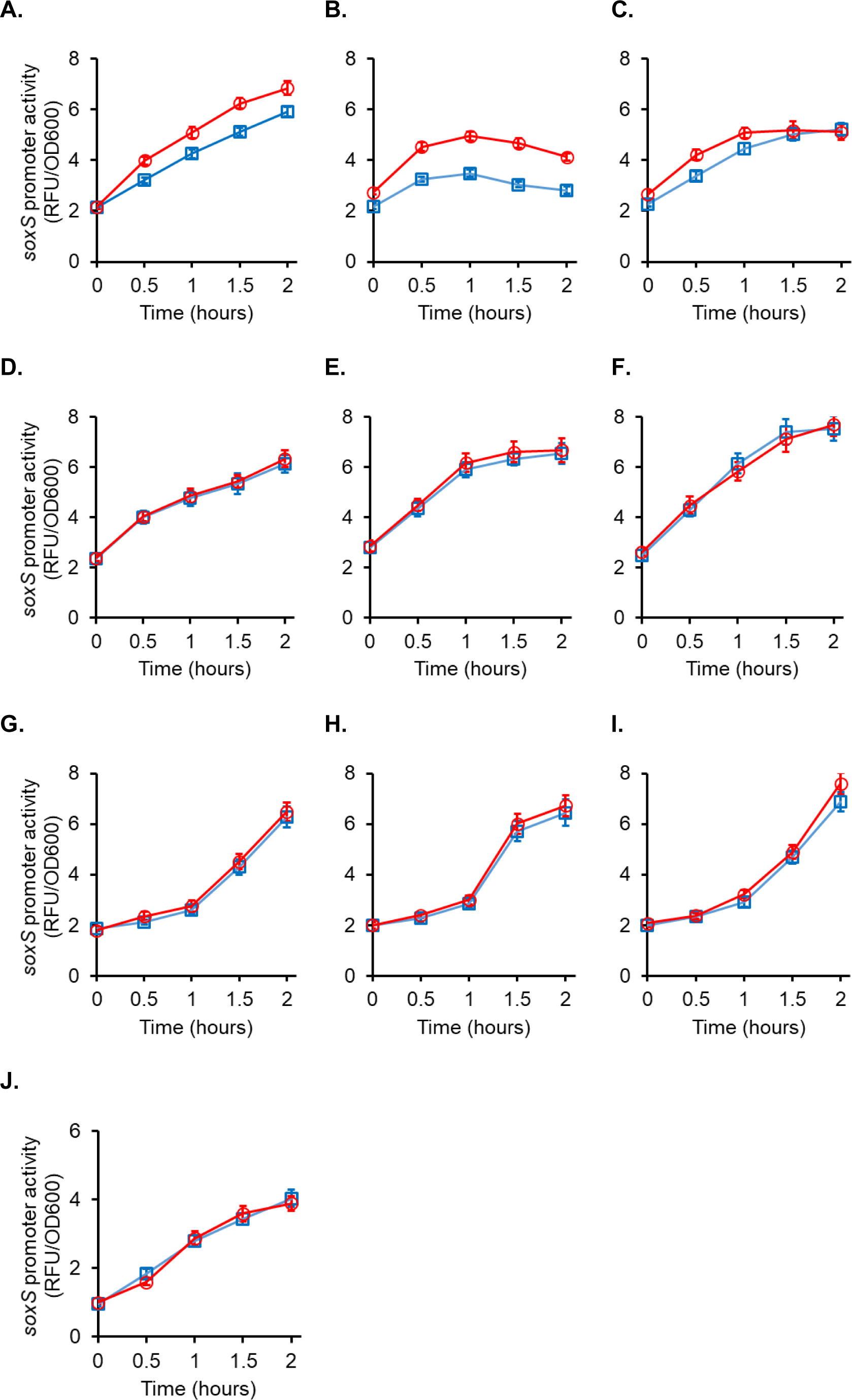
In the presence of paraquat, lines Alt1 **(A)**, Alt2 **(B)**, and Alt3 **(C)** exhibited a faster induction of the *soxS* promoter when brought from M9 glycerol media containing rhamnose (red) compared to cells brought from M9 glycerol media (blue). The rhamnose-dependent faster induction of the *soxS* promoter is absent in the paraquat-evolved lines (PQ1 **(D)**, PQ2 **(E)**, and PQ3 **(F)**) and in rhamnose-evolved lines (Rha1 **(G)**, Rha2 **(H)**, and Rha3 **(I)**). All experiments were done in triplicate. The average of the three experiments and the standard deviation is reported. **(J)** Ancestor cells, when transitioned to M9 glycerol with paraquat from M9 glycerol media with (red) or without rhamnose (blue) do not exhibit any statistically significant difference in the induction kinetics from the *soxS* promoter.

Put together, this data demonstrates that in the lines evolved in alternating rhamnose and paraquat, anticipatory regulation evolves. As we show in the manuscript, this anticipatory regulation leads to a growth phenotype. Our results here demonstrate that one of the manifestations of evolution of anticipatory regulation is partial induction of the *soxS* promoter, when cells are grown in rhamnose. This partial induction leads to a faster transition to the ON state, when cells transition to an environment containing paraquat.

## Discussion

In this work, we present experimental evidence of evolution of anticipatory regulation in bacteria. We demonstrate that by evolving *E. coli* in alternating environments of rhamnose and PQ, anticipatory regulation evolves, where prior exposure to rhamnose enhances fitness of the population in PQ. In our experimental set-up this adaptation evolves in the time scale of a few hundred generations. Anticipatory regulation does not evolve in the first 600 generations, and only evolves thereafter. We show that the first mutations that fix in the evolved environment are the ones which directly enhance fitness. Only when these adaptive mutations are fixed in the population, do mutations that confer adaptive benefit because of anticipatory regulation fix in the population. While the molecular targets which confer advantage because of anticipatory regulation are unknown, the observed fitness advantage is likely because of reprogramming of transcription networks, which evolve faster than other biological networks^87^. This can happen via various events and processes such as gene duplication, horizontal gene transfer, and via point mutations in the transcription factor and/or target promoter^88–90^.

The ecological forces leading to this anticipatory evolution of crosstalk are less clear. Molecular mechanisms for evolution of this phenomenon are not known at this point. How does this crosstalk evolve in a more complex, ecological niche? The relative benefits and costs of anticipatory regulation; and the regularity with which the second stimulus follows the first one are critical parameters, which likely dictate the evolution of anticipatory crosstalk. Our work provides a model to study these questions in a greater detail.

One of the cues in our study is oxidative stress. This was chosen since modelling suggested that anticipatory regulation could evolve if one of the cues caused a growth defect in the bacterium. But this also poses a challenge regarding the specificity of the response. Bacteria and yeast have previously been reported to elicit a general stress response, that is independent of the specific details of the stressor applied ^91–94^. Thus, studying evolution of anticipatory regulation between two stresses poses a challenge. In such an experiment, where S1 and S2 are both stresses, disentangling the effects of cross-regulation and anticipatory behaviour becomes a challenge. When this was attempted in yeast, cross-regulation or anticipatory regulation was observed in a period of 300 generations. However, the authors in such a study were unable to clearly resolve these two effects ^95^. Thus, we argue that one of the two signals should not be associated with a stress. While our modelling results indicate that one of the signals have a strong growth phenotype.

Sequencing parts of genome reveal little details of the molecular mechanisms of this adaptation. This is not surprising. The molecular details of the physiological changes that we characterize likely involve a large number of genes, and are complex in nature. In fact, previous work with yeast indicated has indicated that resistance to a particular stress could be acquired via a number of distinct molecular pathways ^96^. This multiplicity of solutions to an environmental challenge has been predicted by theory too ^97^. In the case of stress, resistance has also been shown to be dependent on the environment that cells were exposed to, prior to introduction of the stress ^96^. Interestingly, stress response in yeast has also been shown to exhibit anticipatory regulation, consistent with how the organism encounters these stresses in its niche ^98^. However, how gene regulation ensuring this evolves remains unknown. Our study is an attempt to answer these questions about evolution of anticipatory regulation.

## Acknowledgements

AM is supported by the Council of Scientific and Industrial Research (CSIR), Government of India, as a Senior Research Fellow (09/087(0873)/2017-EMR-I).

